# Bangers and cash: Baiting efficiency in a heterogeneous population

**DOI:** 10.1101/379743

**Authors:** Naomi Indigo, James Smith, Jonathan K. Webb, Ben L. Phillips

## Abstract

- ***Context***: The uptake of baits is a key variable in management actions aimed at the vaccination, training, or control of many vertebrate species. Increasingly, however, it is appreciated that individuals of the target species vary in their likelihood of taking baits. To optimise a baiting program, then, we require knowledge, not only on the rate of bait uptake, and how this changes with bait availability, but also knowledge on the proportion of the target population that will take a bait.
- The invasive cane toad (*Rhinella marina*) is a major threat to northern quolls (*Dasyurus hallucatus*), which are poisoned when they attack this novel toxic prey item. Conditioned taste aversion baits (cane toad sausages) can be delivered in the field to train individual northern quolls to avoid toads.
- ***Methods:*** Here we report on a large-scale field trial across eleven sites across one large property in Western Australia. Camera trapping and statistical modelling was used to estimate the proportion of baitable animals in the population, and the proportion of these that were baited at varying bait availabilities.
- ***Results:*** Population estimates varied at each site from 3.5 (±0.76 SD) to 18 (± 1.58 SD) individual quolls per site, resulting in a range across sites of 0.6-4 baits available per individual. Bait uptake increased with increasing bait availability.
- We also estimate that only 62% of individual quolls are baitable, and that a baiting rate of 3 baits per individual (rather than per area) will result in almost all of these baitable individuals being treated.
- We compared our statistical method with prior data informing the probability of being baitable; and with probability of being baitable set to 1; this resulted in largely differing estimates in relation to an appropriate baiting rate.
- **Synthesis and applications:** Data and models such as ours provide wildlife managers with information critical to informed decision making and are fundamental to estimate the cost-efficiency of any baiting campaign.

## Introduction

Globally, many populations of wildlife are intensively managed, and the delivery of baits is an important management tool in this context (Bomford & O’Brien 1995). The uptake of doses of toxic or non-toxic compounds in baits is a necessity in the vaccination (Henning *et al.* 2017), training (Gentle, Massei & Saunders 2004; Cagnacci *et al.* 2005) control (Kirkpatrick & Turner 1985) and eradication (Moseby *et al.* 2011; Dundas, Adams & Fleming 2014; Johnston *et al.* 2014; Kimball *et al.* 2016) of many vertebrate species.

A successful baiting program is one that is cost-effective and results in a large proportion of target individuals taking bait (Thomson 1986). Significant progress has been made with the technical aspects of delivering a baiting program, such that baiting can typically be achieved inexpensively and without complex tools or training (Avery et al. 1995). Aircraft, for example, can provide an efficient and fast means of distributing baits over large or otherwise inaccessible tracts of country (Thomson 1986). Success is mostly hampered by the attractiveness and palatability of baits or the willingness of individuals in the population to consume baits. It is increasingly appreciated, for example, that some individuals may be less bait-susceptible, because they are neophobic, bait-shy, or otherwise uninterested when their usual diet is abundant (Birch 1999; Francis *et al.* 2003; Mappes, Marples & Endler 2005; Kelly & Phillips 2017).

Estimating the heterogeneity across a population in the propensity to take baits is a useful first step in assessing a baiting program. A useful second step is to optimise bait delivery by determining the fewest baits required to achieve a given proportion of the population baited. These aims require data on bait uptake within the target population and how this changes with density of baits delivered into the landscape. Collection of these data is often logistically difficult and costly. As a consequence, management decisions are often made based on operator experience rather than empirical evidence (Cook, Hockings & Carter 2010). Remote monitoring tools (such as camera trapping) offer a cost-effective means to acquire the relevant data, but these data do come with analysis challenges. We seek to estimate key parameters (such as the proportion of baitable individuals, and the effect of bait density on uptake probability) from mark-recapture data acquired from camera traps.

Cane toads were introduced in north-eastern Australia in 1935 and have since rapidly expanded across the north of Australia (Phillips *et al.* 2007). The toads carry with them a suite of defensive toxins - - Bufadienolides--unlike toxins possessed by native Australian animals. As a result many vertebrate predators, including northern quolls *(Dasyurus hallucatus),* die after attacking or consuming toads (Covacevich & Archer 1975; Webb, Shine & Christian 2005; Smith & Phillips 2006; Hayes *et al.* 2009; Shine 2010; Webb, Pearson & Shine 2011). Northern quolls are now listed as federally endangered as a consequence of the toad invasion. The delivery of conditioned-taste-aversion baits (cane toad sausages) can, however, be used to train individual northern quolls *(Dasyurus hallucatus)* to avoid toxic invasive cane toads *(Rhinella marina)* (Indigo *et al.* 2018). Conditioned-taste-aversion (CTA) is a powerful innate response that has evolved as a defence mechanism against poisoning (Sinclair & Bird 1984; Conover 1995; Cohn & MacPhail 1996; Mappes, Marples & Endler 2005; Page & Ryan 2005; Glendinning 2007) and results in an animal acquiring an aversion to a referent food as a result of a nauseating or unpleasant experience (Gustavson, Nicolaus & H. Frank 1987). There is currently intense interest in training native Australian predators to avoid cane toads (O’Donnell, Webb & Shine 2010; Webb *et al.* 2015; Ward-Fear *et al.* 2016; Jolly *et al.* 2017; Kelly & Phillips 2017; Ward-Fear *et al.* 2017). Previous research by Indigo *et al.* (2018) shows that quolls consuming a cane toad sausage reduce their attack behaviour towards, and overall interest in, cane toads. Importantly, bait uptake by quolls was also observed when baits were delivered under a realistic field scenario.

Conditioned taste aversion baiting, on a landscape scale is a relatively new technology. Thus the importance of developing evidence-based predictions to guide decision makers is imperative (Jackson *et al.* 2007). But in many ways, it is simply a new baiting technology, so methods developed for assessing CTA baits apply equally to any other bait. In the present paper, we develop an analysis which exploits camera-trap data to estimate the proportion of baitable individuals and how uptake probability changes with bait availability.

## Methods

### Cane toad sausages

Cane toad sausages were made of 15g of minced skinned adult cane toad legs, 1 whole cane toad metamorph (weighing <2g), and 0.06g of Thiabendazole (per sausage; dose rate less than 300mg/kg adult quoll body weight, determined by the smallest – 200g – adult seen at our study site), all packed into a synthetic sausage skin and deployed fresh (Indigo *et al.* 2018). Thiabendazole is an inexpensive, broad-spectrum anthelmintic and antifungal agent (Robinson, Stoerk & Graessle 1965). It is orally-effective and regarded as relatively safe, producing low mammalian mortality: oral LD50 is 2.7g/kg body weight (Dilov *et al.* 1981). Thiabendazole induces a robust CTA after a single oral dose (Nachman & Ashe 1973; O’Donnell, Webb & Shine 2010) and is physically stable at ambient conditions in the bait substrate (Gill, Whiterow & Cowan 2000; Massei, Lyon & Cowan 2003).

### Study Area

The study was conducted in the Artesian Range section (c. 170,000 ha) of Charnley River–Artesian Range Wildlife Sanctuary (16°24’S, 125° 30’E, 300,000 ha), a property managed by the Australian Wildlife Conservancy (AWC) in the Kimberley region of Western Australia. We worked at eleven sites on the property (Fig. 1); sites were selected based on the detection of quolls in the AWC’s fauna surveys (AWC unpub. data, 2017). Each site was separated by at least 5 km to maximise independence between sites. At the time of the study, toads were yet to arrive at any of our sites.

**Figure 1:**
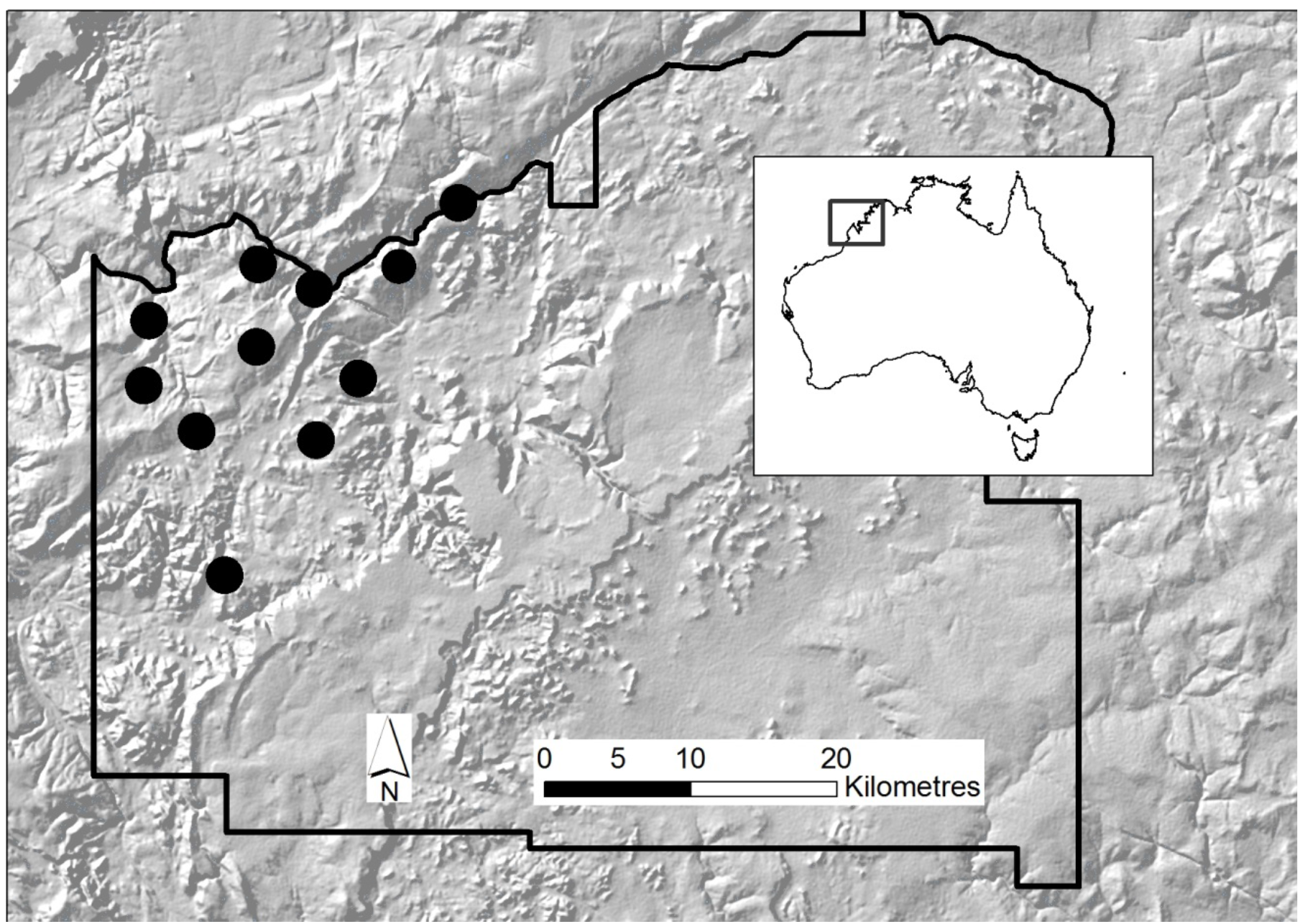
Location of the eleven sites and broader study area within Australian Wildlife Conservancy’s Charnley River – Artesian Range Wildlife Sanctuary, in the Kimberley region, Western Australia.

### Bait stations

In this study, “site” is the location where we deployed an array of bait stations and cameras. “Bait station” is a location within a site where bait was offered. A “session”, is a time interval when bait stations were active. A total of one session was conducted over the course of the seven day period. Each session recorded two “bait events”, where new bait was placed unsecured at a bait station and the old bait removed. In this study, there were two baiting events - BE1 and BE2.

Each site contained 12 bait stations placed 100-120 m apart in a 4 × 3 grid plot array. A single camera trap (White flash and Infrared Reconyx Motion Activated, (HP800, U.S.A) was placed at each bait station. In the first bait event (BE1), the bait was peanut butter (Kraft, Australia) and mackerel in brine (Homebrand, Australia Ltd, Australia) (Hohnen *et al.* 2013; Austin *et al.*2017), available for four nights. In the second bait event (BE2), cane toad sausage baits were made available at the bait station for three nights.

Cameras were secured to trees or rocky ledges approximately 1 m from the ground and aligned to face directly downwards (Hohnen *et al.* 2013; Diete *et al.* 2016). Cameras were set to take five consecutive photographs for each trigger with no delay between triggers. Each bait was placed inside a ring of powdered insecticide (Coopex) to protect from ant spoilage. A total of 132 individual cane toad sausages were deployed across the 11 sites over the period of study.

### Camera trap data collation

Images from bait stations were collated and tagged by pass, site, species, and activity. A ‘pass’ was recorded when a new species entered the frame or when the same species entered the frame following no activity for at least 5 minutes. “Activity” was hierarchical, with the highest activity being ‘Bait taken’; this was defined as either photographic evidence of animal eating bait or bait being taken from the bait station. ‘Bait investigated’ was defined as when bait was sniffed but not consumed or taken. ‘Bait area investigated with no bait available’ was scored when no bait was available at a bait station, but the animal was still visiting or investigating the bait station.

We analysed data using two levels of observation to determine 1) which individuals (and species) were attracted to bait, and 2) which individuals (and species) took bait. We identified individual *D. hallucatus* that visited bait stations by their unique spot patterns (Hohnen *et al.* 2013) to determine visitation rate and bait uptake of individuals. To do this we employed Wild ID (Version 1.0, January 2011) (Bolger *et al.* 2011). We also conducted manual checks with all photographs and compared them to those already identified to determine whether a new individual had been recorded. Quolls were identified to individual within each site, and we treat each site (separated by a minimum of 5km) as independent with regard to quoll ID and behaviour.

### Statistical analysis

#### The model

We assume that there are two classes of individuals: those that will take a bait given the opportunity, and those that will not. We denote the number of individuals at site s, *N_s_.* We are interested in estimating the proportion of *Ns* that are bait-susceptible, z, and the proportion of these baitable individuals, *b_s_,* that are baited under our baiting regime at each site. Our observations consist of a sighting history for each observed individual over seven nights of camera-trapping at the site, and the number of individuals at a site observed to take a bait.

To estimate *Ns* we used a closed population mark-recapture analysis in which each individual (denoted i) was either observed, or not, according to a draw from a Bernoulli distribution:

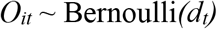

Previous experience with quolls shows that detection rate of individuals declines within seven days (Smith *et al.* 2017), thus we assumed that detection probability (*d_t_* driven by attraction to baited stations) declines over time within each trapping session (Otis *et al.* 1978). We assumed that all individuals had an equal initial detection probability, and that this declined over time according to:

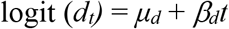

Were *d_t_* denotes the expected detection probability at time *t, μ_d_* is the expected detection probability at t=0,

*B_d_=* change in detection over time.

We used the “data augmentation” method (Royle, Dorazio & Link 2006; Royle, Dorazio & Link 2007; Kery & Schaub 2011) in combination with this detection probability to estimate *N_s_* for each site. Data augmentation offered a flexible way to model patterns of detection probability in our closed populations (Kery & Schaub 2011). Under this approach, the data are ‘padded’ by adding an arbitrary number of zero-only encounter histories of ‘potential’ unobserved individuals. The augmented dataset is modelled as a zero-inflated version of the complete-data model (Royle, Dorazio & Link 2006; Royle, Dorazio & Link 2007) and changes the problem from estimating a count, to estimating a proportion. This was executed by adding a latent binary indicator variable, *R_i_*, to classify each row in the augmented data matrix as a ‘real’ individual or not, where *R_i_* ~ Bernoulli(*Ω*_*s*_). The parameter *Ω*_*s*_ is estimated from the data, and *N_s_* = Σ_*i*(*s*)_ *R_is_*.

Our second observation type was the number of individual quolls in the population observed to take bait on one occasion or more, *T_s_*. We assumed this variable:

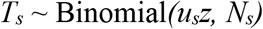

Where *u_s_* is the probability of a quoll at site s taking a bait, given it is a baitable individual. This formulation assumes that uptake probabilities are independent for each individual within a site within our observation window. We make *u_s_* a function of the baits per animal at a site: 12/*N^s^*:

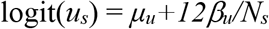

The model was fitted using Bayesian Markov Chain Monte Carlo (MCMC) methods within the package JAGS (Plummer 2016) using Program R (R Core Team 2017). We used minimally informative priors (see Table 1) except for *z*. For this parameter we had prior information on the proportion of quolls likely to take a bait from Indigo *et al.* (2018). Parameter estimates were based on 100,000 iterations with a thinning interval of 5 following a 10,000 sample burn-in. Three MCMC chains were run, and model convergence assessed by eye, and using the Gelman-Rubin diagnostic (Gelman & Rubin 1992b; Gelman & Rubin 1992a).

**Table 1:** Model parameters and their priors including prior distributions, standard deviation, estimated posterior means and their 95% credible intervals. † denotes empty cells. N denotes normal probability distribution, U denotes uniform probability distribution.

| Model Parameters |  |  |  |  |
| --- | --- | --- | --- | --- |
| Name for parameter | Parameter | Distribution (Priors, SD) | Posterior mean | 95% CI |
| Probability of being baitable | $z$ | N (0.63, 0.064) | 0.62 | 0.49, 0.74 |
| Intercept for detection (logit) | $\mu_d$ | N (0,1000) | 1.90 | 0.45, 4.30 |
| Slope of time effect on detection (logit) | $\beta_d$ | N (0,1000) | -0.23 | -0.31, -0.16 |
| Intercept for bait uptake (logit) | $\mu_\mu$ | N (0,1000) | -3.19 | -5.80, -1.32 |
| Slope of bait available effect on uptake (logit) | $\beta_\mu$ | N (0,1000) | 0.39 | 0.32, 0.47 |
| Proportion of augmented set (out of 10,000) | $\Omega$ [1] | U (0,1) | (0,1) | † |
| Proportion of augmented set (out of 10,000) | $\Omega$ [2] | U (0,1) | (0,1) | † |
| Proportion of augmented set (out of 10,000) | $\Omega$ [3] | U (0,1) | (0,1) | † |
| Proportion of augmented set (out of 10,000) | $\Omega$ [4] | U (0,1) | (0,1) | † |
| Proportion of augmented set (out of 10,000) | $\Omega$ [5] | U (0,1) | (0,1) | † |
| Proportion of augmented set (out of 10,000) | $\Omega$ [6] | U (0,1) | (0,1) | † |
| Proportion of augmented set (out of 10,000) | $\Omega$ [7] | U (0,1) | (0,1) | † |
| Proportion of augmented set (out of 10,000) | $\Omega$ [8] | U (0,1) | (0,1) | † |
| Proportion of augmented set (out of 10,000) | $\Omega$ [9] | U (0,1) | (0,1) | † |
| Proportion of augmented set (out of 10,000) | $\Omega$ [10] | U (0,1) | (0,1) | † |
| Proportion of augmented set (out of 10,000) | $\Omega$ [11] | U (0,1) | (0,1) | † |

## Results

### Individual quolls that visited bait stations and bait uptake

Cameras at bait stations detected a total of 86 individual quolls across our 11 sites. During BE2 (in which sausages were deployed), bait stations were visited by 45 individual quolls. Of these 45 individuals, 21 encountered a cane toad sausage with the remaining 24 animals arriving at the bait station after baits had been taken by other species or other quolls. Of these 21 CTA bait-exposed individuals, 18 individuals took the bait (Table 2).

**Table 2:**
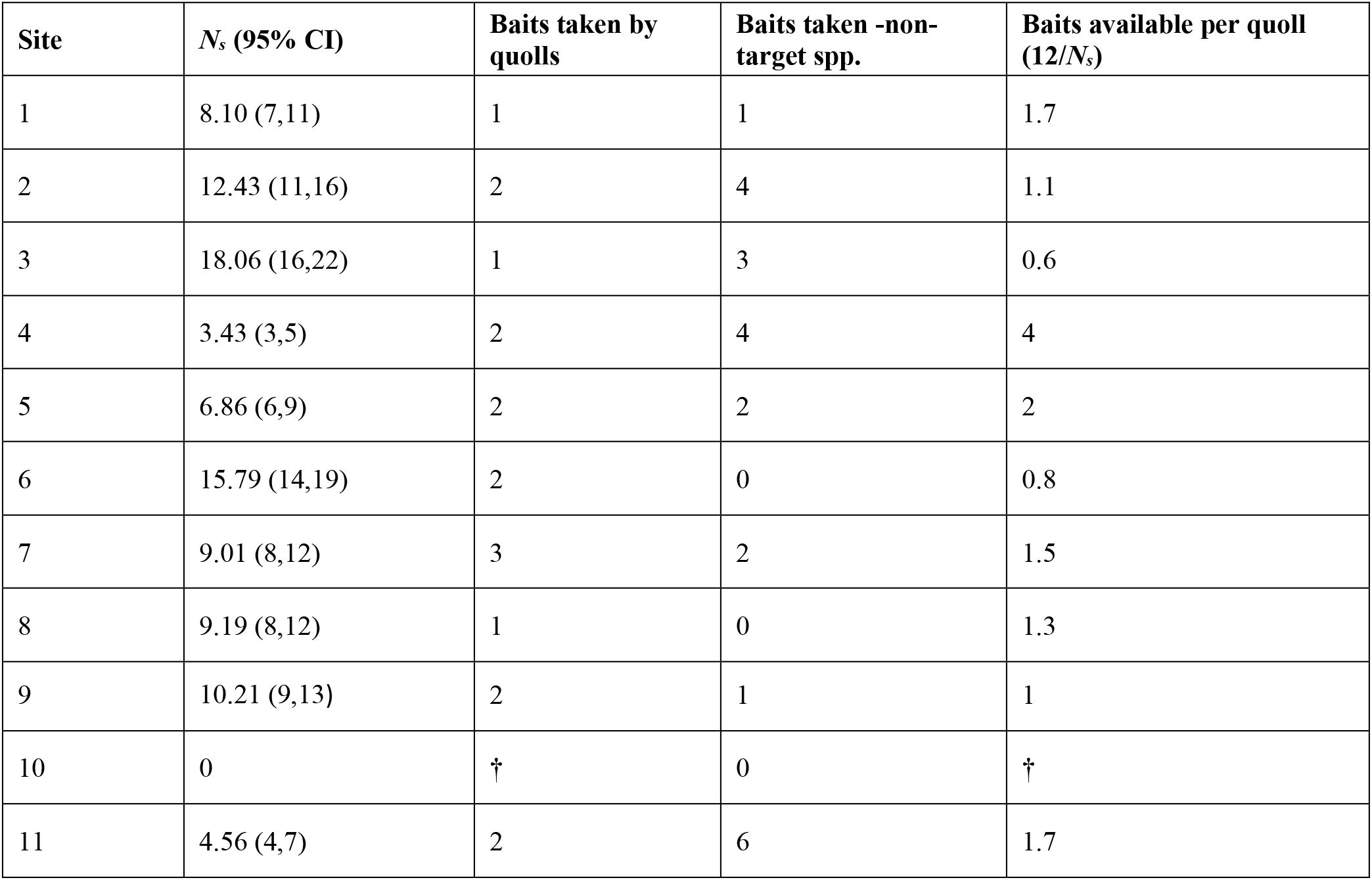
Posterior mean population sizes (*N_s_* (Site 1: Site 11)) and 95% credible intervals, assuming closure of the population during the time of the study. Number of baits taken by quolls at each site, number of baits available per quoll at each site was a function of *N_s_* because there were 12 baits deployed at each site. † denotes empty cells.

### Non-target CTA bait uptake

A total of 23 species were identified investigating bait stations. Of these non-target species, bait was taken by *Corvus orru* = 6, *Zyzomys argurus* = 4, *Isoodon macrourus* = 4, *Ctenotus spp.* = 3, *Mesembriomys macrurus* = 1, Unknown rodent sp. = 1.

### The model

Bayesian mark-recapture population estimates (*N_s_*) and associated credibility intervals (CI = 95%) for each site are listed in Table 2. Baiting at the nominal rate of 1 bait per 100m^2^ resulted in 0.63-4 baits available per animal (Table 2). There was a clear positive relationship between the probability of bait uptake and number of baits available per animal at each site (β_μ_ =0.40; 95% CI: 0.32 – 0.47).

Baiting at one bait per 100m^2^ resulted in bait uptake by around 18% of quolls at a site. The proportion of baitable quolls *z* = 0.62 (95% CI: 0.4911 - 0.7417). Baiting at the rate of 3-5 baits per individual should result in essentially all baitable individuals taking bait (Fig.2).

**Figure 2:**
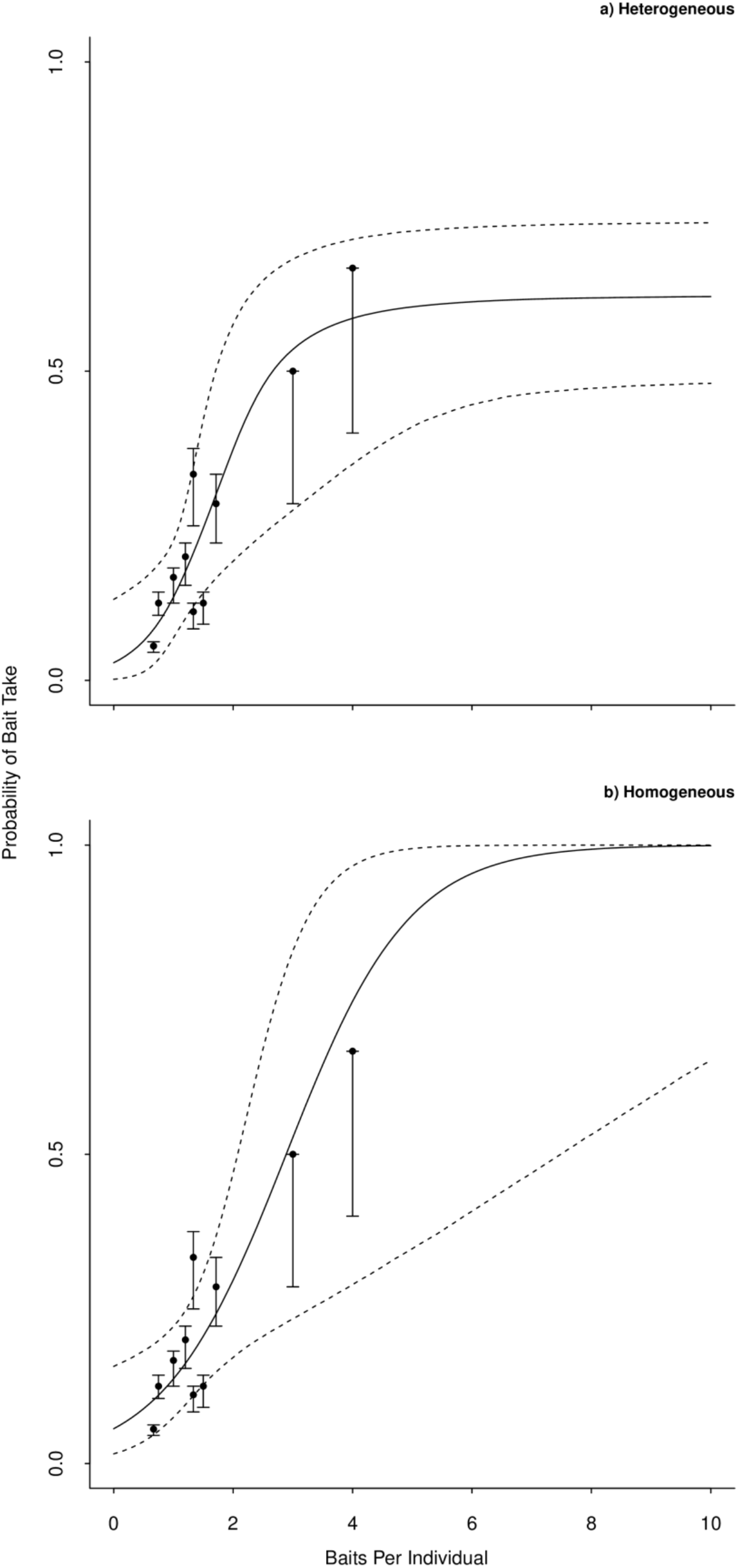
Modelled fitted logistic curve with the raw data overlayed (black circle and associated prediction envelope), dashed line denotes 95% prediction envelope. Comparison of fitted curve with (a) and without (b) updated prior *z*. Plot a) Heterogeneous represents suitable baiting rate when *z=0.62,* baiting rate to saturate all baitable individuals = 3-5 baits per individual animal, plot b) Homogeneous plot represents baiting rate when z=1.0, baiting rate to saturate all baitable individuals = 7-8 baits per individual.

## Discussion

Here we have used standardised camera trap arrays across 11 sites to generate data on bait uptake, and how this varies with bait availability. We used these data (with some prior information) to estimate key parameters of interest: the proportion of a population that is baitable, and how the uptake probability increases with bait availability. These parameter estimates provide valuable information for managers considering a baiting program, allowing them to assess the likely outcome of a baiting program, optimise the baiting rate, and estimate costs. In our case, the baiting program is to train wild quolls to avoid cane toads immediately prior to the arrival of cane toads in the landscape.

We estimate that the best outcome achievable in our system is for us to deliver baits to 62% of the population. The other 38% of the population appear to not be baitable. Most of our information on this parameter comes from our prior expectation of uptake rate, derived from a combination of field and captive trials by Indigo *et al.* (2018). We updated our prior value for *z* – the variable representing the probability of being baitable; with these data, to obtain the posterior value of 62%. We observed only a small shift (Fig. 3); thus our new observations simply provide greater certainty on this parameter, indicating that the value of 62% baitable animals is robust across two field sites, and animals from two different origins (WA and NT). Such variation in baitability may be consistent with general observations of variation in food preference within populations (Birch 1999). Alternatively we might be observing variation in boldness: some individuals may exhibit more of a neophobic response to bait as a consequence of genetic predisposition (Marples *et al.* 2007; Hoppitt & Laland 2013; Greggor *et al.* 2014). These predispositions might have developed as a result of early learning, experience, interactions with the environment and the feeding preferences of adults (Birch 1999; Francis *et al.* 2003; Mappes, Marples & Endler 2005; Hoppitt & Laland 2013).

**Figure 3:**
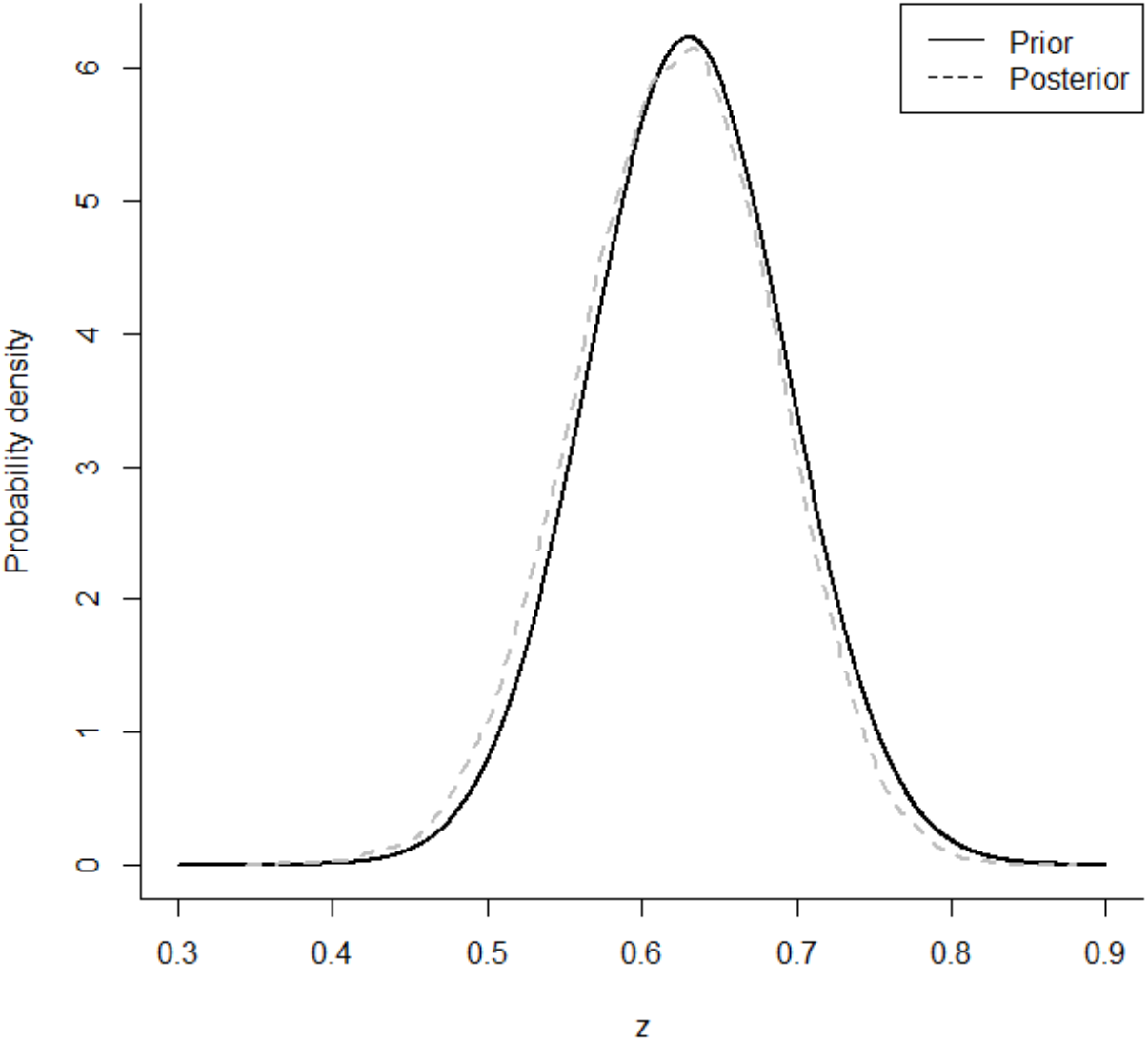
Posterior and prior distribution of *z* (probability of being baitable). Prior distribution (solid black line) and posterior (dashed grey line) estimates the proportion of a population that is baitable. Posterior for *z* is shifted slightly when we updated this prior.

Irrespective of the mechanism, non-baitable individuals reduce the proportion of the population that are trainable with CTA. While 62% may seem a relatively poor level of bait success, we can reasonably expect that some fraction of these non-baitable animals may also avoid toads. Certainly there is evidence in other taxa that some level of innate avoidance of toads is present in toad-naïve predator populations (e.g., the common planigale (*Planigale maculata*) (Webb *et al.* 2008), red-cheeked dunnart (*Sminthopsis virginiae*) (Webb, Pearson & Shine 2011), native rodents (*Melomys burtoni, Rattus colletti* and *Rattus tunneyi*) (Cabrera-Guzmán *et al.* 2015), terrestrial snake species (Phillips, Brown & Shine 2003) scincid, agamid and varanid lizards (Smith & Phillips 2006; Llewelyn *et al.* 2014; Pearson *et al.* 2014), avian species (Beckmann & Shine 2009), and aquatic animals such as the barramundi (*Lates calcarifer*), sooty grunter (*Hephaestus fuliginosus*) (Crossland 2001) northern trout gudgeons, (*Mogurnda mogurnda),* Dahl’s aquatic frogs (*Litoria dahlia*) (Nelson, Crossland & Shine 2011) and freshwater crocodiles (*Crocodylus johnstoni*) (Somaweera & Shine 2012). There is also evidence that such innate avoidance may provide the raw material on which natural selection can act to generate a rapid adaptive response to toads (Phillips & Shine 2006; Smith & Phillips 2006; Llewelyn *et al.* 2011; Somaweera & Shine 2012; Ujvari, Oakwood & Madsen 2013; Kelly & Phillips 2017; Tingley *et al.* 2017). Clearly, then, this non-baitable fraction of the population are important, but the fraction of these non-baitable animals that will in fact avoid toads is unknown and remains an important avenue for future work.

Our analysis also gives us insight into an optimal baiting rate. We found that bait uptake increases with increasing bait availability. This is to be expected, and has also been demonstrated in lethal baiting programs (Christensen, Ward & Sims 2013). Interestingly, some studies suggest that low bait availability can reveal further population heterogeneity; in foxes (*Vulpes vulpes*), it is the dominant individuals that access baits first, consequently reducing access to baits for other individuals within the population (Marks & Bloomfield 1999; Gentle, Massei & Saunders 2004). Our findings suggest that sub-optimal baiting rates can be avoided in our case by increasing baiting density to 3 baits per individual; resulting in a very high probability of bait take by the baitable fraction of northern quolls.

Bait availability is defined against the number of animals at a site, but our analysis also gives us insight into the density of quolls and how this varies within the landscape. Baiting at a rate of one bait per hectare in the northern Kimberley, and assuming an average population density, managers may reasonably expect bait uptake from an estimated 18% of the northern quoll population. Why such a low percentage? At a density of one bait per hectare, many quolls are simply not encountering the bait; arriving at the bait station after baits have been taken. Deploying between 3-4 baits per hectare, however, can be expected to treat almost all of the baitable quolls at a site. Our results might also imply that baits could be applied at one per hectare on three different occasions, but this strategy was not actually one we tested, and may result in poorer uptake probability than a simple three-fold increase in bait density.

Another reason to not bait on three separate occasions is cost. A substantial cost associated with our baiting program is helicopter time. Three separate baiting occasions would result in a threefold increase in helicopter cost. On the other hand, increasing bait density would generate only a moderate increase in cost (if any). Increasing baiting density three-fold does, however, generate a three-fold increase in bait production cost; a cost increase whether baits are delivered on one occasion, or three. Alternatively, for a fixed budget, increasing bait density reduces the area that can be treated by a factor of three. Clearly, then, given a fixed budget, the optimal baiting strategy in our case will depend upon this trade-off: are we better off to treat a larger area (but fewer animals per area), or a smaller area (but treat almost all the baitable animals in that area)? This is not a trivial problem to solve, and will require application of spatially explicit population viability analysis. Nonetheless, our finding should sound a note of caution with regard to deployment strategies of CTA baits. Training only once prior to toad arrival will need to be delicately timed: too early, and trained animals may lose their aversion before toads arrive (Indigo *et al.* 2018). Delivering baits over three occasions may increase the probability of animals receiving training prior to the arrival of cane toads. The precision of such timing is complicated by inevitable uncertainty with regard to where the toad invasion front is, and when it will arrive at the site. Thus, any baiting campaign will need to dedicate effort to predicting the date of toad arrival at the site.

Our results provide important information for designing a baiting program for quolls in the Kimberley. While our results may be used to guide programs elsewhere, this should only be done with caution. Many important variables change between areas. For example, bait uptake may change with patterns of seasonal variation in diet (Oakwood 1997), reproductive status of individuals (Oakwood, Bradley & Cockburn 2001), the suitability and complexity of the target species habitat (Hohnen *et al.* 2016a; Hohnen *et al.* 2016b); the availability of preferred/alternative prey (Algar *et al.* 2007; Weerakoon & Banks 2011; Christensen, Ward & Sims 2013) and of course,the extent to which non-target species access baits before the target species. Our results are nonetheless encouraging with regard to the use of toad sausages as a vehicle for large-scale CTA training of quolls. We might only expect to treat around 62% of animals, but we can treat this proportion of the population using a feasible baiting rate: three sausages per individual. The ability to predict the results of CTA baiting provides wildlife managers greater flexibility in decision making. From the perspective of developing a working baiting strategy, our results contribute to a growing body of work opening the possibility for broad-scale CTA quoll training using toad aversion sausages: a technique to prevent quolls from attacking cane toads (O’Donnell, Webb & Shine 2010; Webb *et al.* 2015; Jolly *et al.* 2017; Kelly & Phillips 2017; Indigo *et al.* 2018).

More broadly, our results speak to the importance of assessing population heterogeneity in baitability. There is clear evidence that individuals vary in their food preference and behavioural tendency to accept novel food. If our analysis had not accounted for this heterogeneity, and assumed that all individuals were baitable, we may well have suggested that all individuals could be treated if only we increased our bait density 2-fold (Fig. 2). By accepting that there is heterogeneity, we do not fall into this trap, and instead, observe how increasing bait availability results in saturating all baitable animals. As a consequence, we are able to design a more efficient baiting program. We achieved this inference using data obtainable from camera trap arrays which is also encouraging. It is increasingly feasible to gather the data required informing the design of baiting programs, and models such as ours allow us to capture the key parameters of interest to managers and decision makers.

## Authors’ Contributions

All persons who meet authorship criteria are listed as authors, and all authors certify that they have participated sufficiently in the work to take public responsibility for the content, including participation in the concept, design, analysis, writing, or revision of the manuscript.

## Acknowledgements

We thank Australian Wildlife Conservancy and their supporters for access to their field laboratory and for contributing to the study. We wish to also thank Katherine Tuft and Sarah Legge for their efforts during the preliminary stages of project. Many thanks to Kristina Koenig, whose efforts assisted with individual identification of quolls. This project was supported by The Holsworth Wildlife Research Endowment & The Ecological Society of Australia. Additionally we wish to thank all who participated with field trials.

## Compliance with Ethical Standards

The study was conducted under Wildlife Conservation Regulation 17 (Permit number: SFO10584), and approved by the University of Technology Sydney Animal Care and Ethics Committee (Protocol: 2012-432A) and Department of Parks and Wildlife Animal Ethics Committee (Protocol: AEC 2016_50 and Protocol 2013_37). Additionally this study was conducted in accordance with the approved outline submitted to the AVPMA, Permit number: PER92262.

